# Tunability of calcium dynamics by signaling inputs and cell-cell communication in pancreatic beta cells

**DOI:** 10.1101/2024.10.31.621335

**Authors:** Xingbo Shang, Andre Levchenko

## Abstract

In addition to frequently occurring stable all-or-none responses, live cells can display more complex response dynamics, e.g., oscillations in the activity/concentration of biomolecules. While the emergence and function of oscillatory dynamics have been heavily investigated, fewer efforts have been spent on whether cells can fine-tune different aspects of an oscillatory signal (e.g., peakwidth, duty cycle, and frequency) and whether this fine-tuning can allow oscillatory signals to convey different information. In this study, we investigate glucose-induced calcium (Ca^2+^) oscillation in MIN6 cells, finding that the spontaneous or induced changes in the phosphatidylinositol 4,5-bisphosphate (PIP2) level during Ca^2+^ oscillation can modulate the peakwidth of individual Ca^2+^ spikes. Additionally, using a combination of optogenetics and Ca^2+^ imaging, we demonstrate that variation of the widths and frequencies of Ca^2+^ spikes can exert strong influence on coupling between neighboring MIN6 cells.

## Introduction

The pancreatic beta cell, the workhorse of glucose homeostasis, utilizes intracellular calcium (Ca^2+^) as a key second messenger to relay an increase in extracellular glucose level to insulin secretion. Cells are first depolarized by the closing of the ATP-sensitive potassium channel (KATP) due to the glucose-dependent increase in ATP production. The depolarization then leads to opening of voltage dependent calcium channel (VDCC), allowing extracellular Ca^2+^ to rapidly flow into the cell. The influx of Ca^2+^ further depolarizes the cell and triggers multiple negative feedbacks including but not limited to 1) increased ATP consumption which releases its blockage of KATP, 2) opening of the voltage dependent potassium channels (Kv) leading to the plasma membrane repolarization and 3) increased extrusion of Ca^2+^ from the cytoplasm by proteins such as the sodium calcium exchanger (NCX) or sarcoendoplasmic reticulum Ca^2+^-ATPase (SERCA).^1,2^ This complex regulatory mechanism commonly triggers intracellular Ca^2+^ oscillations upon glucose stimulation, thought to be at least partly responsible for the pulsatile secretion of insulin.^3,4^

Due to its close linkage to insulin secretion, malfunctioning of this Ca^2+^ circuit in beta cells can disturb glucose homeostasis. For example, activating mutations in KATP can cause neonatal diabetes^5,6^, while loss-of-function (LOF) mutations of KATP can lead to hyperinsulinism^7,8^. Mutations in other glucose related proteins (e.g. glucose transporter^9^ and glucokinase^10^) have also been reported to lead to diabetes. Recently, it was found that patients with long QT syndrome, a disease caused by LOF mutations of KCNQ1 (Kv7.1) or KCNH2 (Kv11.1), also suffer from higher burden of diabetes.^11^ Additionally, mutations in KCNQ1 have been identified by multiple studies to be associated with diabetes.^12-14^ Unlike other proteins with well established roles in glucose sensing, Kv7.1 and Kv11.1 have received considerably less attention in beta cell analysis, possibly because of the observations that Kv2.1 is the major Kv for repolarizing beta cell.^15,16^ Therefore, why malfunctioning of these two ‘minor’ Kv channels can increase risk of diabetes remains largely unknown.

Complex intracellular Ca^2+^ dynamics and its alterations can have important functional consequences. From a quantitative point of view, when compared with a sustained increase in Ca^2+^, oscillatory Ca^2+^ dynamics offers more tunable parameters: frequency, duty cycle, peak shape, etc. Beta cells can indeed show a diversity of oscillation frequencies emerging not only in single cells but also in the ensembles of coupled cells: a study by Nunemaker et al., found that islets of Langerhans isolated from different mice can have significant heterogeneity in the oscillation period, ranging from ∼0.5min for faster ones to ∼5min for slower ones.^17^ Additionally, the Ca^2+^ dynamics observed from a single islet under constant glucose stimulation can also switch between different patterns.^18,19^ Over the past decades, extensive efforts have been dedicated to identifying new proteins or small molecules involved in the generation of Ca^2+^ signals^20-22^ and developing a quantitative understanding of how oscillation can emerge from the interactions of these biomolecules^23-25^. However, these studies usually focused on the assaying of either presence or absence of Ca^2+^ oscillation in beta cells with less attention paid to how cells can diversify the Ca^2+^ signals by fine-tuning the oscillation after it is turned on.

Another intriguing question about Ca^2+^ in beta cells is the emergence of synchronized Ca^2+^ dynamics when beta cells exist in the form of 3D islet. Such synchronization was disrupted when beta cells were dispersed onto 2D surface^26^ or connexin-36, a gap junction protein, was knocked out^27^, suggesting the importance of cell-cell communication in synchronization. Two recent studies provide strong evidence that a subgroup of super-connected beta cells exist in islet and can act as pacemakers to generate synchronized Ca^2+^ dynamics.^28,29^ These observations suggest that instead of generating its own Ca^2+^ dynamics, a beta cell may also follow the pace set by its neighbors through cell-cell communication, but it remains unclear how cell-cell coupling influences calcium dynamics quantitively or even qualitatively. For example, it is not clear if single beta cells can discriminate and prioritize the heterogeneous inputs it receives from multiple neighboring beta cells.

In this study, we addressed these two challenges in the quantitative understanding of beta cell calcium signaling by investigating the origin of diversity in Ca^2+^ dynamics and its role in cell-cell communication in a beta cell model cell line, MIN6. By combining Ca^2+^ imaging with pharmacological perturbations, real-time optogenetic manipulations, and mathematical modeling, we found that changes in phosphatidylinositol 4,5-bisphosphate (PIP2), associated with hormonal control of pancreatic cells, can enhance the diversity of Ca^2+^ dynamics by fine-tuning the oscillations. Additionally, depending on the relative activity of Kv7 and Kv11.1, the outcome of PIP2-dependent effects can be different. Finally, using optogenetic stimulation, we demonstrate that during cell-cell communication, transient spiking, when compared with long-lasting plateau, can be more effective in coupling to neighboring cells.

## Results

### Calcium oscillations relying on Kv7 or Kv11.1 ion channels show differential dynamical behavior

To explore the roles of the Kv7 and Kv11.1 channels in glucose-induced Ca^2+^ dynamics, cells were first preincubated with saturating concentrations of Dofetilide (Kv11.1 inhibitor) and XE991 (Kv7 inhibitor) before glucose stimulation to inhibit both Kv11.1 and Kv7 channels and Ca^2+^ dynamics were recorded from individual cells with GCaMP3^30^. After observing that most cells (85 out of 95) adapted to a Ca^2+^ plateau under this condition, the concentration of either Dofetilide or XE991 was lowered until ∼50% of cells showed oscillatory Ca^2+^ dynamics (Fig. 1A). For convenience, we have referred to the cells with oscillatory Ca^2+^ dynamics due to reduced inhibition on Kv7 or Kv11.1 as ‘the Kv7 oscillator’ or ‘the Kv11 oscillator’ respectively. Interestingly, it was observed that in addition to constant oscillation, Ca^2+^ levels in a portion of cells can undergo single or multiple switching between oscillation and non-oscillatory plateau levels (Fig. 1B and 1C).

**Figure 1.**
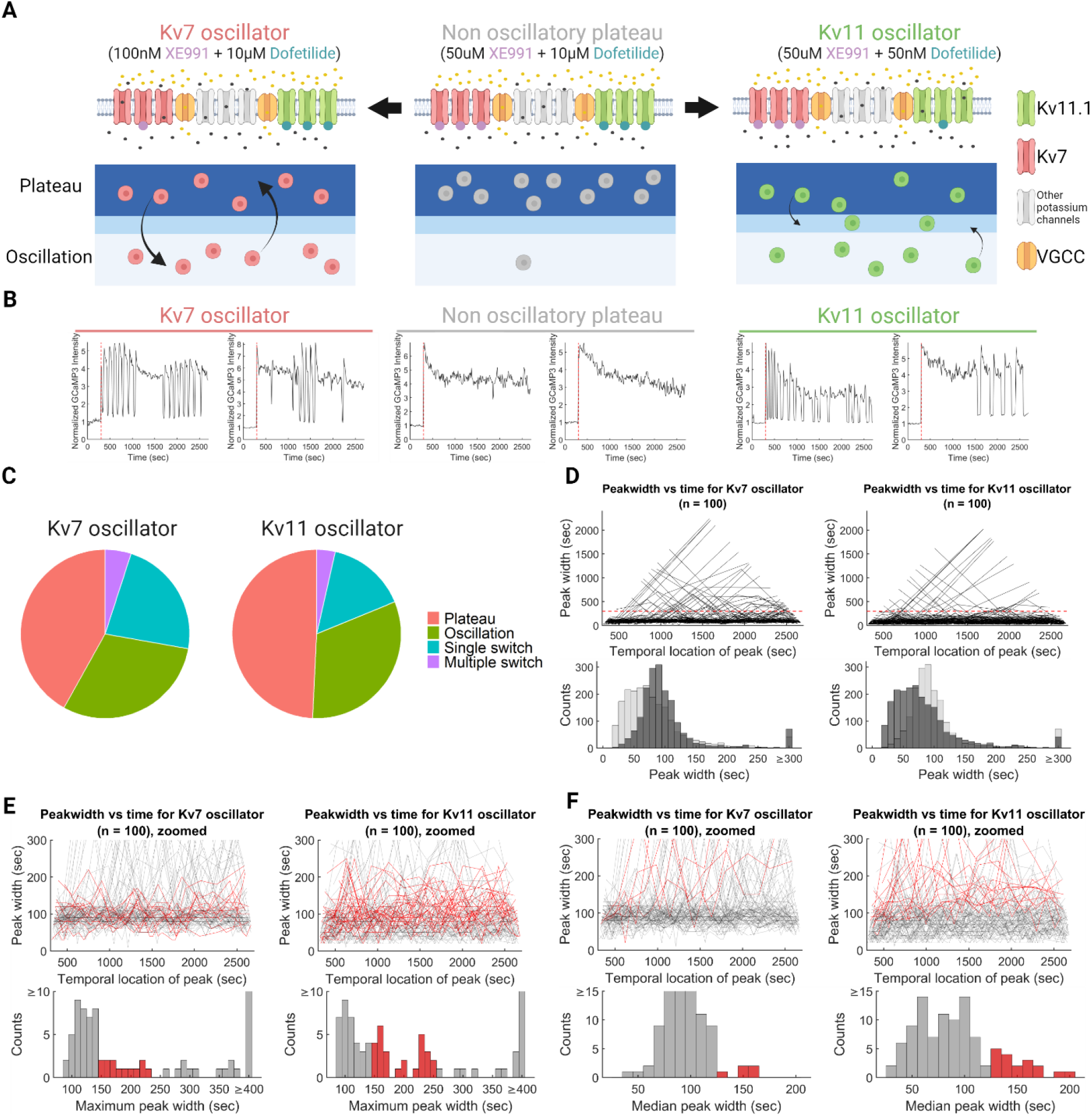
A) Schematic illustration of Kv7 and Kv11 Ca^2+^ oscillator and their corresponding dynamical behavior of Ca^2+^. B) Representative Ca^2+^ dynamics of multiple switching and plateau from Kv7 or Kv11 oscillator and non-oscillatory cells. C) Overall statistics of number of cells with different types of Ca^2+^ dynamics from Kv7 and Kv11 oscillators. D) Time series of peakwidth (upper panel) and overlapped histograms of peakwidth of all Ca^2+^ peaks (lower panel) for Kv7 (left panel) and Kv11 oscillators (right panel). In the overlapped histogram, bars of the corresponding oscillator are highlighted in dark grey. E) Zoomed view of peakwidth time series in D) (upper panel) and histograms of maximum Ca^2+^ peakwidth for each cell (lower panel). Time series with maximum peakwidth in the range of 150∼250sec are highlighted in red. Bars within the same range are highlighted in red in the histograms. F) Zoomed view of peakwidth time series in D) (upper panel) and histograms of median Ca^2+^ peakwidth for each cell (lower panel). Time series with median Ca^2+^ peakwidth in the range of 130∼200sec are highlighted in red. Bars within the same range are highlighted in red in the histograms.

To more systematically explore the difference between the Kv7 and Kv11 oscillators, the width of each peak (the ‘peakwidth’) in the Ca^2+^ dynamics was quantified to generate time series of the change in the peakwidth over time for each cell. As shown in Fig. 1D, when compared with Kv11 oscillator, Kv7 oscillator had a more concentrated distribution of peakwidth and a higher tendency to switch from oscillation to plateau (a plateau was operationally defined as a “peak” with the peakwidth≥300sec). Additionally, the more populated intermediate range of maximum peakwidth (Fig. 1E) and the longer tailing in distribution of median peakwidth (Fig. 1F) further demonstrated Kv11 oscillator’s tendency to maintain oscillation without abrupt change in peakwidth when compared with Kv7 oscillator.

### Kv7 and Kv11 oscillators show high sensitivity to PIP2 perturbation with different responses

We then explored possible mechanisms underlying the differences between Kv7 and Kv11 oscillators. Previous research has shown that the conductance of Kv7 has a stronger dependency on phosphatidylinositol 4,5-biphosphate (PIP2) than Kv11.1.^31^ We thus hypothesized that the change in peakwidth is due to a decrease in Kv conductance and that Kv7 oscillator changes more abruptly due to the higher sensitivity of Kv7 conductance to PIP2.

To test this hypothesis, GSKA1^32^, a specific inhibitor of phosphatidylinositol 4-kinase type IIIα (PI4KA), was used to block PIP2 production following the initiation of oscillation generated by Kv7 and Kv11 oscillators. When added at saturating concentration (10nM), GSKA1 almost completely silenced Ca^2+^ signals from both Kv7 and Kv11 oscillators. This can be due to a known effect of PIP2 depletion on VDCC.^33,34^ To observe if any change in peakwidth can be induced by GSKA1, the dosage of GSKA1 was titrated, and it was found that at 2 nM, two types of the switch in Ca^2+^ dynamics can be induced: to damped oscillation or adapted plateau (Fig. 2A.). More specifically, we classified the response as the damped oscillation, if Ca^2+^ level maintained oscillation with slight increase/decrease in peakwdith and a gradually damped amplitude. By contrast, during adapted plateau, Ca^2+^ output initially reached a level higher than the baseline and then gradually decreased. In agreement with our hypothesis, we consistently found that the Kv7 oscillator showed a stronger tendency to switch to adapted plateau (an abrupt change in the peakwidth) while Kv11 oscillator tended to switch to damped oscillation (with no or mild change in the peakwidth), following addition of 2 nM of GSKA1 (Fig. 2B).

**Figure 2.**
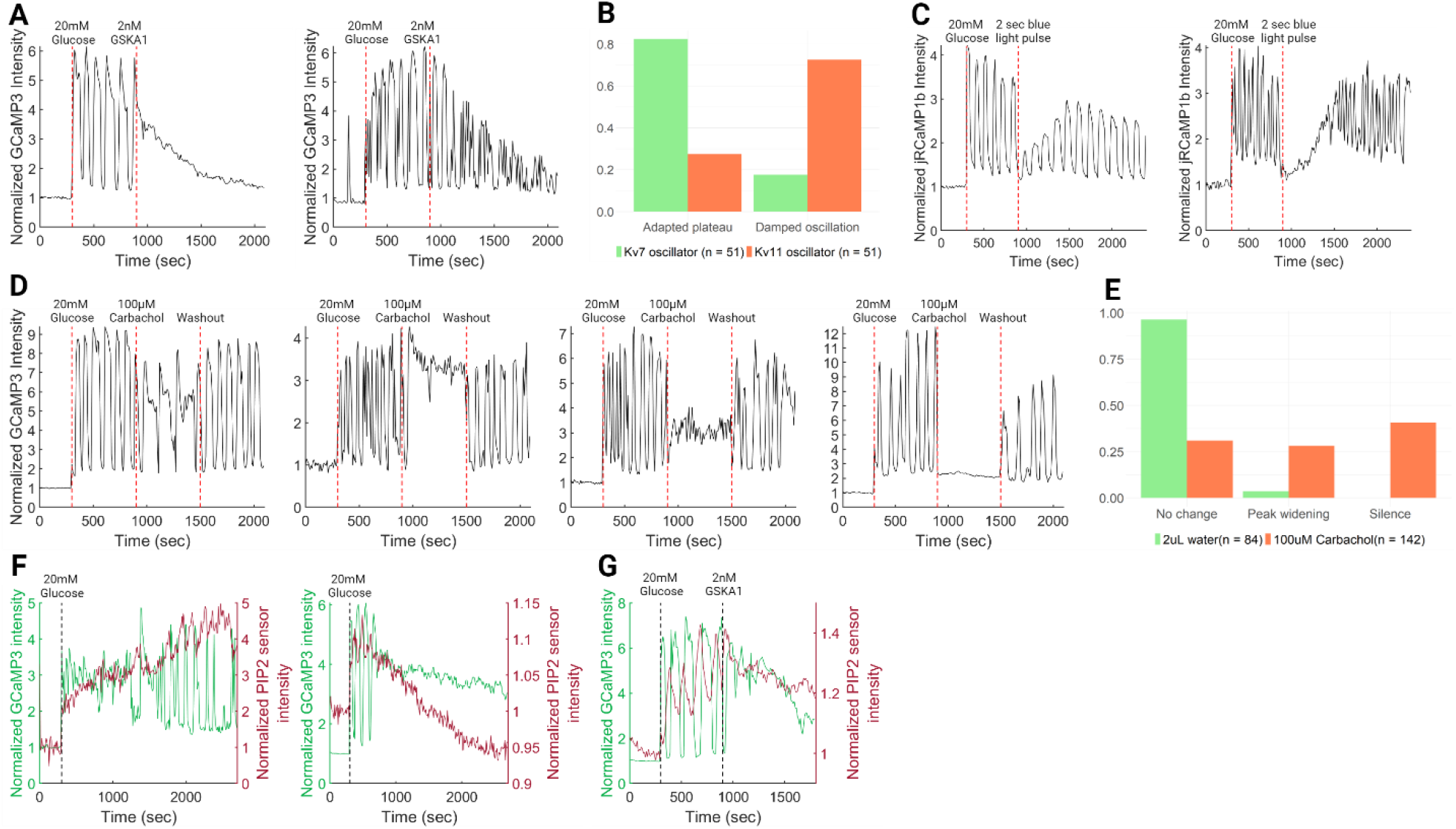
A) Representative Ca^2+^ dynamics of adapted plateau (left panel) and damped oscillation (right panel) generated by addition of 2nM GSKA1 10min after glucose-induced Ca^2+^ oscillation. B) Percentage of cells showing adapted plateau or damped oscillation for Kv7 and Kv11 oscillators after addition of 2nM GSKA1. C) Representative Ca^2+^ dynamics generated by blue light induced PIP2 consumption by CRY2-OCRL and CIBN-Caax. D) Representative Ca^2+^ dynamics generated by addition of 100μM Carbachol 10 min after glucose-induced Ca^2+^ oscillation and washout of carbachol after 10min incubation. E) Percentage of cells with no change, peak widening or silence for cells treated as in D). For negative control, an equal volume of water, the solvent of carbachol, was added in place of carbachol. F) Representative Ca^2+^ and PIP2 dynamics of cell switching from plateau to oscillation (left panel) or from oscillation to plateau (right panel) in Ca^2+^ dynamics. G) Representative Ca^2+^ and PIP2 dynamics of cell switching from plateau to oscillation in Ca^2+^ dynamics after addition of 2nM GSKA1.

To rule out the possibility that our observations of damped oscillation and adapted plateau were caused by nonspecific effects of GSKA1, an orthogonal approach based on blue light-inducible PIP2 consumption^35^ was applied to Kv7 oscillator in combination with the red fluorescent Ca^2+^ sensor jRCaMP1b^36^. As shown in Fig. 2C, upon a two-second pulse of blue light, Ca^2+^ oscillation was abruptly silenced due to a sudden loss of PIP2 and gradually recovered back to an oscillatory response in two distinct regimes: through either an oscillation or plateau with a gradually increasing amplitude. Thus both GSKA1 inhibition and direct decrease of PIP2 abundance, lead to a rapid damping or adaptation of the Ca^2+^ response.

In vivo, beta cells can rapidly consume PIP2 through PLC activation when muscarinic acetylcholine receptor M3 (CHRM3) is stimulated by acetylcholine.^37-39^ To demonstrate whether CHRM3 induced PIP2 consumption can shape Ca^2+^ dynamics, CHRM3 was overexpressed in MIN6 cells and 100 μM carbachol was applied to activate CHRM3 10 min after glucose-induced Ca^2+^ oscillations generated by Kv7 oscillator. Upon addition of carbachol, Ca^2+^ spikes underwent significant broadening and dampening, resembling the effect of GSKA1 and light-induced PIP2 consumption, with this effect removed and Ca^2+^ signaling recovered after washout of carbachol (Fig. 2D and 2E).

### PIP2/Ca^2+^co-imaging reveals strong coupling between PIP2 and Ca^2+^ dynamics in Kv7 oscillator

After confirming that PIP2 perturbations can shape Ca^2+^ dynamics of the Kv7 oscillator, we explored whether the spontaneous changes of PIP2 levels during Ca^2+^ signaling can interface with Ca^2+^ dynamics. To achieve this, we combined a recently described red fluorescent PIP2 sensor^40^ with the green fluorescent genetically encoded Ca^2+^ biosensor, GCaMP3, to co-image PIP2 and Ca^2+^ dynamics in Kv7 oscillator.

This co-imaging revealed in cells showing stable oscillation that PIP2 also displayed oscillatory dynamics, and that Ca^2+^ oscillated in an out-of-phase manner, such that the peak of Ca^2+^ coincided with the trough of PIP2 and vice versa (Fig. S1). This observation agrees with previous publications of PIP2/ Ca^2+^ co-imaging and may be explained by Ca^2+^ induced activation of PLC, thus generating a Ca^2+^-PLC-PIP2 feedback loop.^41,42^ We then focused on cells undergoing a switch between oscillation and plateau and found a strong correlation with the change in PIP2 signal amplitude: 8 out of 9 cells spontaneously switching from plateau to oscillation in Ca^2+^ dynamics showed an increase in PIP2 level while 14 out of 17 cells switching from oscillation to plateau in Ca^2+^ dynamics showed a decrease in PIP2 level (Fig. 2F). Moreover, the decrease in PIP2 level and switch from Ca^2+^ oscillation to plateau induced by GSKA1 resembled the PIP2/ Ca^2+^ dynamics recorded from cells undergoing spontaneous switching between these two Ca^2+^ signaling regimes (Fig. 2G).

### Mathematical modeling accounts for the roles of PIP2 and Kv channels in shaping calcium dynamics

To unify experimental observations, we modified a previously published mathematical model^24^. We first explored the role of ATP-and voltage-sensitive potassium channels (KATP and Kv, respectively) in shaping the Ca^2+^ dynamics. Indeed, although the focus of our experimental analysis has been on the roles of different Kv channels, KATP might also influence the observed Ca^2+^ dynamic. The model suggested that, at low Kv and KATP conductance values (upper left dark purple region of Fig. 3A), Ca^2+^ levels stabilized at a constantly high plateau, as the low conductance through these channels cannot repolarize the cell. Similarly, at high Kv and KATP conductance values (lower right dark purple region of Fig. 3A), Ca^2+^ level stabilized at a constantly low level (silence) as the high conductance prevents depolarization of the cell. When Kv and KATP conductance values are within the range promoting the oscillatory dynamics, decreasing either Kv or KATP conductance leads to an increase in the peakwidth, but varying Kv conductance leads to change of peakwidth over a wider range than varying KATP conductance (Fig. 3A and 3B). In other words, the peakwidth of Ca^2+^ spikes is more sensitive towards the change in Kv conductance.

**Figure 3.**
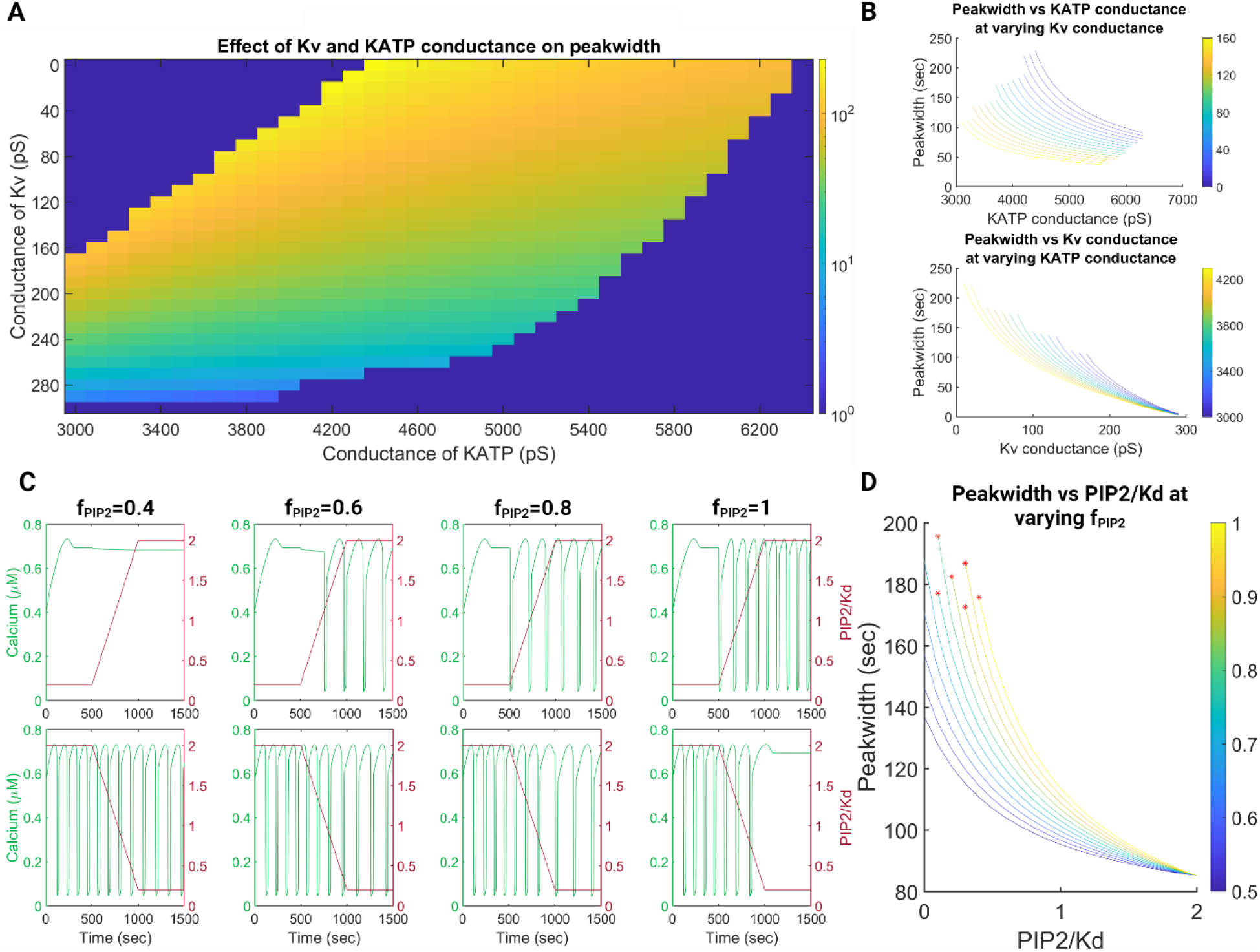
A) Heatmap of peakwidth of Ca^2+^ dynamics simulated at different combinations of Kv and KATP conductance. Values of peakwidth are displayed as color temperature in log scale. The dark purple region in the upper left corner stands for stable plateau Ca^2+^ signal and the dark purple region in the lower right corner stands for silence in Ca^2+^ signal. B) Line graph visualization of A), color temperature of lines correspond to conductance of Kv (upper panel) or KATP (lower panel). C) Simulated response of Ca^2+^ dynamics to an increase (upper panel) or decrease (lower panel) in PIP2 availability (PIP2/Kd) at different *f*_*PIP2*_. D) Effects of PIP2 availability on Ca^2+^ peakwidth simulated at different *f*_*PIP2*_. *f*_*PIP2*_is displayed as color temperature in log scale. The points where Ca^2+^ oscillation switches to plateau are marked by red asterisk signs.

We then introduced PIP2 dependence of Kv conductance into the model. To account for the fact that 1) Kv channels with different PIP2 sensitivities coexist in the same cell and, 2) the exact level of PIP2 dependent conductance for certain types of Kv channels (e.g. Kv11.1^33,43-45^) remains controversial, we introduced PIP2 sensitive fraction(*f*_*PIP*2_), a parameter ranging from 0 to 1 to model the dependence of Kv conductance on PIP2 without losing generalizability. For a specific type of Kv channel, *f*_*PIP2*_ reflects its level of PIP2 dependence. For a cell expressing multiple types of Kv channels, *f*_*PIP2*_ reflects the expression level weighted average of PIP2 dependence of each type of Kv channels. In the scope of this research, higher *f*_*PIP2*_was used to model Kv7 oscillator because Kv7 has higher PIP2 sensitivity than Kv11. To simulate the effect of PIP2 availability without arbitrarily assuming the concentration of PIP2 and dissociation constant of Kv-PIP2 complex (Kd), we define PIP2 availability as the ratio of these two values (PIP2/Kd) and only vary this ratio in our model. Indeed, by varying PIP2 availability at different levels of *f*_*PIP2*_(Fig. 3C and 3D), peakwdith of Ca^2+^ oscillation changed more abruptly at higher *f*_*PIP2*_, reproducing the observations for Kv7 and Kv11 oscillator shown in Fig. 1.

Overall, the combination of experimental analysis and mathematical modeling suggests that PIP2 dependence through Kv7 expression in beta cells can introduce sensitivity to additional external stimuli, such as activation of the muscarinic acetylcholine receptors, but can also make Ca^2+^ signaling more variable in the absence of such signaling, due to the spontaneous fluctuations of PIP2 levels. Furthermore, the dependence on PIP2 can modulate the amplitude and frequency of Ca^2+^ oscillations (reflected in the peakwidth). These findings also raised the question of how the properties of Ca^2+^ signaling in neighboring cells, which can be modulated by spontaneous or induced PIP2 dynamics, might influence communication between these cells and thus the collective cell dynamics essential for coordinated insulin secretion by multiple cells within an islet.

### Optogenetic manipulation of Ca^2+^ dynamics shows separated spikes induce stronger response from neighboring cells than adapted plateau

To understand how cells interpret Ca^2+^ signals with different peakwdith from neighboring cells, we sparsely (MOI ∼0.01) transfected ChR2XXL^46^ fused with Halo tag into MIN6 cells. The transfected cells were then stained by Janelia Fluor 646 Halo tag ligand (JF646) to highlight cells expressing ChR2XXL and Ca^2+^ dynamics was recorded from all cells through Calbryte 630, a Ca^2+^ sensitive dye spectrally orthogonal to both JF646 and ChR2XXL (Fig. 4A). Cells were then stimulated by blue light pulses with two different temporal patterns to induce an adapted plateau lasting for 120 seconds (‘pattern 1’, using multiple closely spaced light pulses with 10sec interval between adjacent pulses) or two separated spikes with 100 seconds interval in between (‘pattern 2’, using two widely spaced light pulses) (Fig. 4B). Hereafter, for convenience, cells expressing ChR2XXL are termed the hub cells (cells with optogenetically induced Ca^2+^ signaling communicating with neighbor cells), and the cells surrounding hub cells (i.e., cells that can be stimulated by the hub cells) are termed the follower cells.

**Figure 4.**
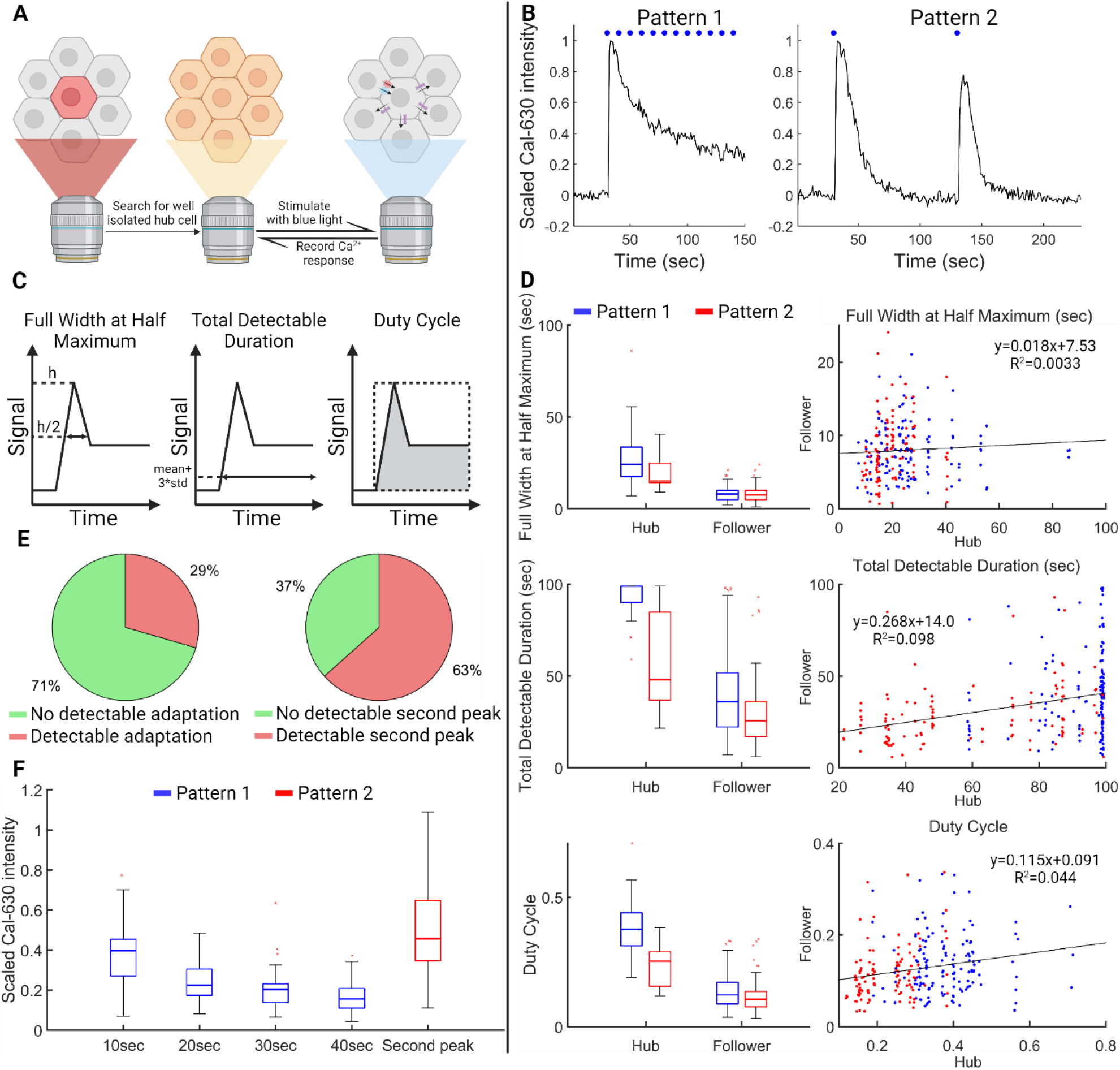
A) Schematic illustration of workflow for simultaneous optogenetic Ca^2+^ manipulation and recording. B) Representative Ca^2+^ dynamics generated by stimulating cells with ‘pattern 1’ and ‘pattern 2’. C) Schematic illustration of the definition of FWHM, total detectable duration, and duty cycle. For total detectable duration, signal is considered as detectable if it is above mean plus three times standard deviation of baseline. For duty cycle, areas under the curve are divided by the rectangular areas formed by the peak of Ca^2+^ trajectory and the time after blue light stimulation. D) Comparison of FWHM, total detectable duration and duty cycle between hub and follower cells during the adapted plateau generated by ‘patten 1’ (blue) or the first spike generated by ‘pattern 2’ (red). E) Statistics of followers capable of following the adapted plateau generated by ‘pattern 1’ (left) or the second spike generated by ‘pattern 2’ (right). F) Scaled intensity of followers of adapted plateau generated by ‘pattern 1’ binned at different time points (blue) and followers of second spikes at peak generated by ‘pattern 2’ (blue).

Recording Ca^2+^ dynamics from hub and follower cells revealed that upon blue light stimulation, a Ca^2+^ wave consistently initiated from hub cells and propagated into follower cells (Fig. S2.). We first focused on whether the follower cells could discriminate between an adapted plateau and a transient spike in the hub cells through a differential response. To explore this, we quantitively compared hub and follower cells’ Ca^2+^ dynamics following the ‘pattern 1’ stimulation and the first spike induced by ‘pattern 2’ examining three response measures: full width at half maximum (FWHM), total detectable duration and duty cycle (Fig. 4C). While hub cells stimulated by ‘pattern 1’ had significantly higher values in FWHM, total detectable duration and duty cycle than those stimulated by ‘pattern 2’, the differences were largely abolished in follower cells, especially for FWHM and duty cycle (Fig. 4D). For total detectable duration, we observed that a small subset of follower cells could have long-lasting detectable Ca^2+^ signal above baseline (Fig. 4E). Nevertheless, most follower cells responded with a transient spike regardless of the input from hub cells.

We then compared the capability of adapted plateau (‘pattern 1’) and separated spikes (‘pattern 2’) to induce response beyond one single transient spike in follower cells. For hub cells with adapted plateau, only 29% follower cells could have detectable Ca^2+^ signal (above mean plus three times standard deviation of baseline) lasting longer than 50 sec. By contrast, when hubs cells were stimulated to generate two separated spikes, the second spike could induce detectable response from 63% of the follower cells (Fig. 4E). Additionally, while the Ca^2+^ signal in follower cells adjacent to the hub cells stimulated with ‘pattern 1’ on average rapidly decreased within 40 sec to ∼20% of the peak intensity, the second spike in follower cells adjacent to the ‘pattern 2’-stimulated hub cell on average reached >40% intensity of the first peak in the second spike of the response, with some cells capable of achieving equal or even higher amplitude spike (Fig. 4F).

### Mathematical modeling shows separated spikes induce stronger response than plateau by allowing cells to recover from negative feedback on Ca^2+^

To understand why separated spikes were more effective in triggering response from follower cells than the continuous signaling during the plateau phase, the propagation of Ca^2+^ wave from hub cell to follower cells was modeled as an injection of current into follower cells. As shown in Fig. 5, when a constant current injection was made, the Ca^2+^ level rapidly adapted to a level close to the baseline, similar to the observations from follower cells responding to the ‘pattern 1’ hub cells. This adaptation can be explained by the negative feedback from opening of Kv and KATP channels and deactivation of VDCC. When pulsed current injection with increasing intervals in between was made, negative feedbacks had sufficient time to recover to baseline and repetitive Ca^2+^ spikes similar to follower cells of ‘pattern 2’ hub cells could be generated.

**Figure 5.**
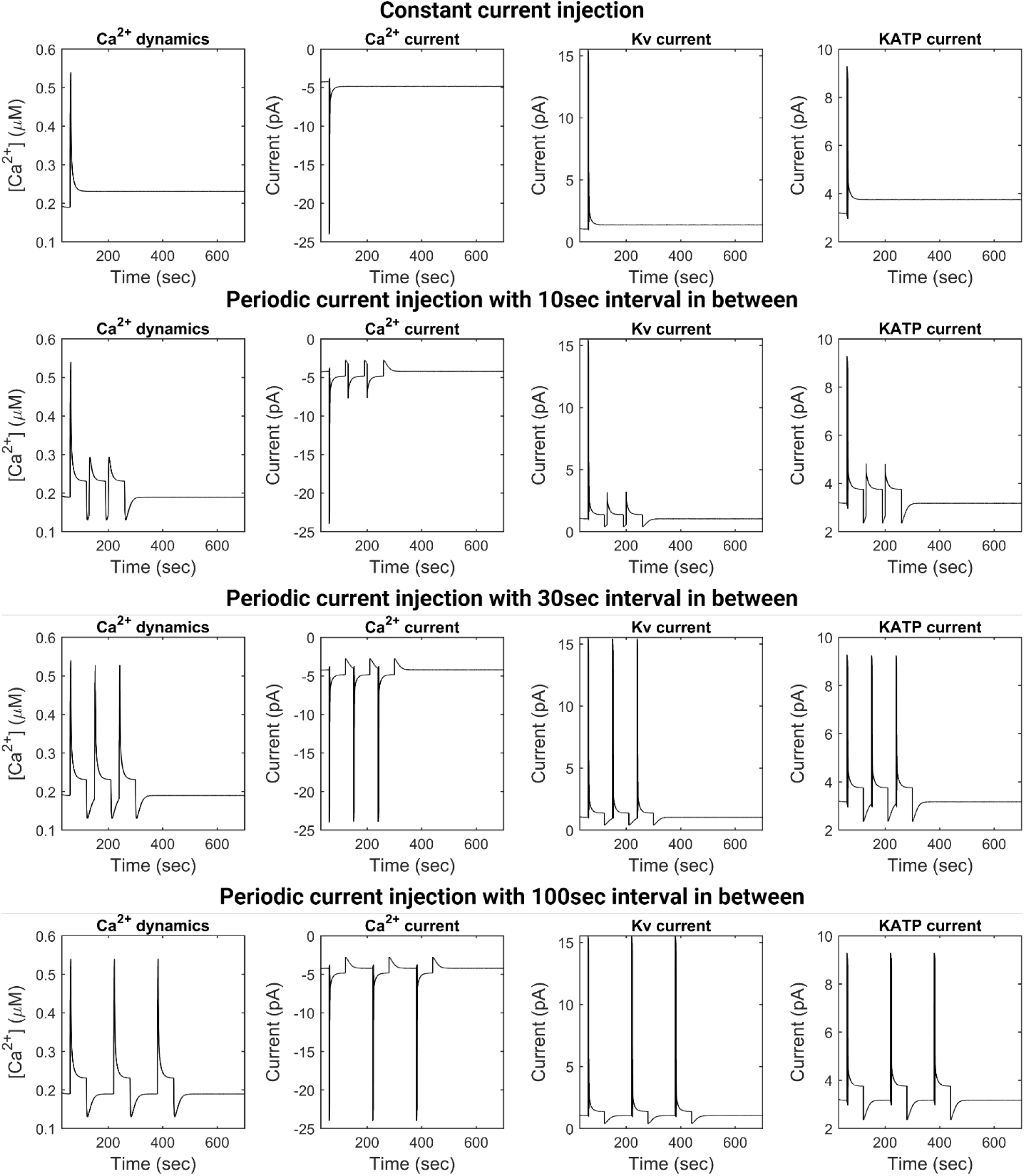
Simulated Ca^2+^ dynamics, Ca^2+^ current, Kv current and KATP current in response to constant current injection or periodic current injection of -0.3pA lasting for 60sec with 10sec, 30sec, 100sec of interval between two adjacent injections.

## Discussion

Oscillatory signals can provide multiple ways of encoding information in the signaling dynamics, which can provide a much richer signal encoding-decoding opportunities. Changes in parameter values controlling the oscillatory responses can also permit switching from dynamically variable to non-oscillatory, plateau signals. Therefore, it is theoretically possible for cells to diversify oscillatory dynamics by only varying the activity/concentration of biomolecules already existing in the circuit without rewiring.

In this study, we explored this theoretical possibility in the context of Ca^2+^ signaling in beta cells. We find that if two seemingly loosely related features: 1) high sensitivity of Ca^2+^ peakwidth to Kv conductance and 2) different types of Kv channels differ in PIP2 sensitivity, coexist in the same molecular circuit, they can allow cells to diversify the width of oscillation peaks in a relatively abrupt (Kv7 oscillator) or continuous (Kv11 oscillator) manner by varying PIP2 level. While similar diversity might also be achieved by heterogeneous gene expression, the mechanism described in this study has multiple unique features. First, it allows cell to swiftly change peakwidth on the timescale of minutes rather than hours required for gene expression. For short term tasks, such as insulin secretion, this short term dynamical responsiveness can be especially important, as it offers the feasibility to rapidly fine-tune insulin secretion. Additionally, by incorporating Kv channels with different sensitivity to PIP2 into the circuit, the responsiveness towards change to the environmental cues controlling PIP2 level (e.g. acetylcholine) can also be fine-tuned. Kv7 oscillator can abruptly change peakwidth upon PIP2 level change and can serve as sensors for such environmental cues. The Kv11 oscillator is less responsive to change in PIP2 level and can serve as a pacemaker to maintain stable oscillation. The co-existence of oscillators with different expression level of Kv7 and Kv11 may allow the whole islet to reach the optimal balance between stability and responsiveness. In other words, the function of Kv7 and Kv11 in beta cells can extend beyond repolarizing cell during Ca^2+^ signaling. This may explain why mutations in Kv7.1 or Kv11.1 can increase the susceptibility to diabetes^11-14^ even though they are not the major carrier of repolarizing current in beta cells^15,16^.

In addition to intracellular signaling, the diversity in Ca^2+^ peakwidth can also have functional role in intercellular communication. By artificially inducing adapted plateau or separated spikes in a subgroup of cells, we found that separated spikes can induce stronger response from the coupled follower cells. This result can also be interpreted in another way: cells can decrease its influence on follower cells by increasing the peakwidth. While this can also be achieved by silencing of Ca^2+^ signal, increasing peakwidth allows other Ca^2+^ dependent process, e.g. insulin secretion, to be less affected.

## Conclusion

This study reveals a new mechanism for diversification of oscillatory Ca^2+^ dynamics where a physiological change in PIP2 level during Ca^2+^ signaling can change the peakwidth of Ca^2+^ spikes in a Kv channel dependent manner. The resulting diversity in peakwidth is further found to have functional role in cell-cell communication as neighbor cells can be better coupled and undergo collective responses when signaling by separated transient peaks rather than isolated long lasting increases in the Ca^2+^ levels. One limitation of this study is that it was carried out in MIN6 cells, a model cell line rather than primary beta cells or islets, and thus may not faithfully reflect the situation in vivo. Nevertheless, the results presented offer the hypothetical mechanisms that can be further tested in more physiological model systems.

## Methods

### Cell culture

MIN6 cells passage 55 to 70 was maintained in DEME supplemented with 15% FBS and 50uM 2-mercaptoethanol. Mycoplasma was routinely checked by DAPI staining.

### Plasmids and lentivirus packaging

Plasmids for GCaMP3 (#22692), CIBN-CAAX (#79574), GFP-CRY2-OCRL (#79565), jRCaMP1b (#63136), and pPHT-PIP2 (#60937) were purchased from Addgene. CHMR3-IRES-mCherry(VB221027-1606ryn) was purchased from VectorBuilder. For constructing ChR2XXL-Halo, fragments encoding ChR2XXL, Halo tag with 2X(GGGGS) linker, and a customized lentiviral backbone with SFFV promoter and WPRE element were PCR amplified by Q5 high fidelity master mix (NEB, M0492S) and assembled by NEBuilder® HiFi DNA Assembly Master Mix (NEB, E2621S).

For lentivirus packaging, the ChR2XXL-Halo plasmid was co-transfected with lentiviral packaging plasmid mix (Cellecta, CPCP-K2A) at a ratio of 2:5 into HEK293FT cell with lipofectamine 3000 (ThermoFisher, L3000008) according to manufacturer’s protocol. Lentivirus was harvested 48 and 72h after transfection, 0.22μm filtered and concentrated 30 times by lenti-X concentrator (Takara Bio, 631231). The concentrated lentivirus was then aliquoted and stored at - 80°C until use.

### Recording unperturbed or GSKA1 perturbed Ca^2+^ dynamics from Kv7 and Kv11 oscillator

To enhance cell adherence, all glass bottom petridish (Matsunami glass, 10810-054) were treated by UV ozone for five minutes and sterilized by UV for 30min before cell seeding. 1.2*10^6^ MIN6 cells were seeded onto UV ozone treated glass bottom petridish and allowed to attach for 24h. Cells were then transfected with 1μg GCaMP3 with lipofectamine 3000 following manufacturer’s instructions. 36∼48h after transfection, cells were washed three times with imaging buffer (125mM NaCl, 5mM KCl, 1mM CaCl_2_, 1mM MgCl_2_, 20mM HEPES, pH adjusted to 7.3∼7.4) and incubated for 1h in a 37°C incubator without CO_2_. Cells were then transferred onto a Zeiss Axiovert widefield microscope equipped with a 37°C incubator and incubated with corresponding concentrations of XE991 and Dofetilide (50μM XE991 + 50nM Dofetilide for Kv11 oscillator; 100nM XE991 + 10μM Dofetilide for Kv7 oscillator) for 30min. At the end of incubation, 5min of baseline Ca^2+^ activity was recorded, and glucose was added into the petridish to a final concentration of 20mM. For unperturbed Ca^2+^ dynamics, cells were then imaged for another 40min. For perturbation with GSKA1, cells were imaged for 10min before corresponding concentrations (2nM or 10nM) of GSKA1 was added. After addition of GSKA1, cells were imaged for another 20min. Unless otherwise stated, all images were acquired with a 10sec interval. A mercury lamp (Excelitas, X-cit exacte) was used as excitation light source. A customized Matlab script was used to extract Ca^2+^ dynamics from raw tiff files. All Ca^2+^ dynamics were background subtracted and then normalized by the mean of baseline signal.

### Perturbation of Kv7 oscillator by light induced PIP2 consumption

Cells were transfected with 1μg jRCaMP1b, 0.5μg CIBN-CAAX and 1μg GFP-CRY2-OCRL and handled similarly for recording unperturbed Ca^2+^ dynamics. After 5min of baseline and addition of 20mM glucose, cells were imaged for 10min and then stimulated by 2sec pulse of blue light and imaged for another 30min.

### Perturbation of Kv7 oscillator with CHRM3

Cells were transfected with 1ug GCaMP3 and 1ug CHRM3-IRES-mCherry and handled in the same way as for recording unperturbed Ca^2+^ dynamics. After 5 min of baseline and addition of 20mM glucose, cells were imaged for 10min. 100μM carbachol was then added to cells and cells were imaged for another 10min. Imaging buffer with carbachol was then replaced with prewarmed imaging buffer containing 20mM glucose, 100nM XE991 and 10uM Dofetilide and cells were imaged for another 10min.

### PIP2 and Ca^2+^ co-imaging of Kv7 oscillator

Cells were transfected with 1ug GCaMP3 and 2ug pPHT-PIP2 and handled and imaged in the same way as for recording unperturbed Ca^2+^ dynamics.

### Mathematical modeling of PIP2, Kv, and KATP’s role in shaping Ca^2+^ dynamics

For the effect of Kv and KATP conductance on peakwdith of Ca^2+^, a published model^24^ was used with differential equations shown below. Physical meanings and values for all parameters are summarized in Table S1.

The change of membrane potential (V) as a consequence of ion current conducted by VDCC (*I*_*Ca*_), Kv (*I*_*Kv*_) and KATP (*I*_*KATP*_) was modeled by:

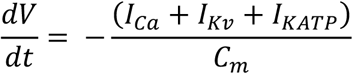

The opening and closing of VDCC, Kv and KATP were modeled by:

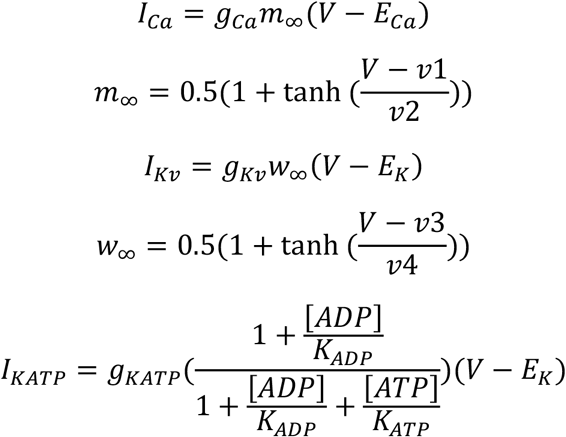

The change of cytoplasmic Ca^2+^ level caused by Ca^2+^ inflow from VDCC and Ca^2+^ extrusion was modeled by:

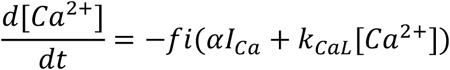

The change of ATP and ADP level through Ca^2+^ dependent metabolism was modeled by:

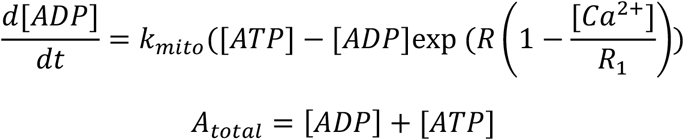

To study the effect of Kv and KATP conductance on peakwidth, g_Kv_ and g_KATP_ were varied from 0 to 300pS with step of 10pS and 3000 to 6400pS with step of 100pS. To introduce PIP2 sensitivity to Kv current, the original equation for modeling Kv current was modified into:

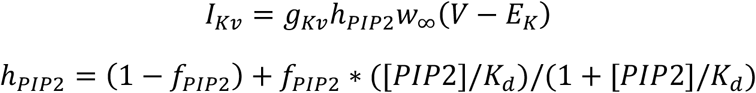

where *f*_*PIP2*_stands for fraction of PIP2 sensitive conductance and [*PIP2*]/*K*_*d*_ stands for PIP2 availability. For Fig. 3C, PIP2 availability was varied linearly from 0.2 to 2 or 2 to 0.2 within 500sec. For Fig. 3D, PIP2 availability was varied from 0 to 2 with a step size of 0.1 and *f*_*PIP2*_was varied from 0.5 to 1 with a step size of 0.05.

### Optogenetics experiment of cell-cell communication

1. *10^6^ MIN6 cells were seeded onto UV ozone treated glass bottom petridish and allowed to attach for 24h. ChR2XXL-Halo was then transfected into MIN6 cells through lentivirus to reach an MOI of ∼0.01. 36∼48h after transfection, 200nM JF646 (Promega, GA1120) and 2μM Calbryte 630 AM (AAT Bioquest, 20720) were added to the culture medium and incubated for 1h. Cells were washed three times with imaging buffer supplemented with 20mM glucose and immediately mounted onto microscope. Well isolated hub cells with ChR2XXL-Halo were identified by fluorescence from JF646. For Ca^2+^ imaging with Calbryte 630, the excitation light was filtered by a 599/13 band pass filter (Chroma, ET599/13x) to minimize activation of ChR2XXL. After recording a 30sec baseline, cells were stimulated by either pattern 1 (twelve 1ms blue light pulses with 10sec interval) or pattern 2 (two 1ms blue light pulses with 100sec interval) with Ca^2+^ imaged simultaneously. All imaging in this section was done with an interval of 1sec. Ca^2+^ dynamics were then extracted, normalized by mean of baseline, and FWHM, total detectable duration and duty cycle were calculated with a customized Matlab script. For calculating total detectable duration, a signal with intensity higher than mean plus three times standard deviation of baseline intensity was treated as detectable.

### Mathematical modeling of cell-cell communication

For modeling cell-cell communication, KATP and Kv conductance was set to 4300pS and 300pS and all other parameters remain unchanged (Table S1). The propagation of Ca^2+^ from hub cell into neighboring cell was then modeled as an extra current (*I*_*hub*_) of -0.3pA added to the total current:

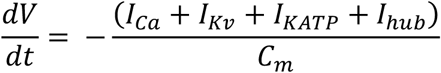

## Notes

### Competing Interest Statement

The authors have declared no competing interest.

