## Supplementray figures and tables for "Tunability of calcium dynamics by signaling inputs and cell-cell communication in pancreatic beta cells"

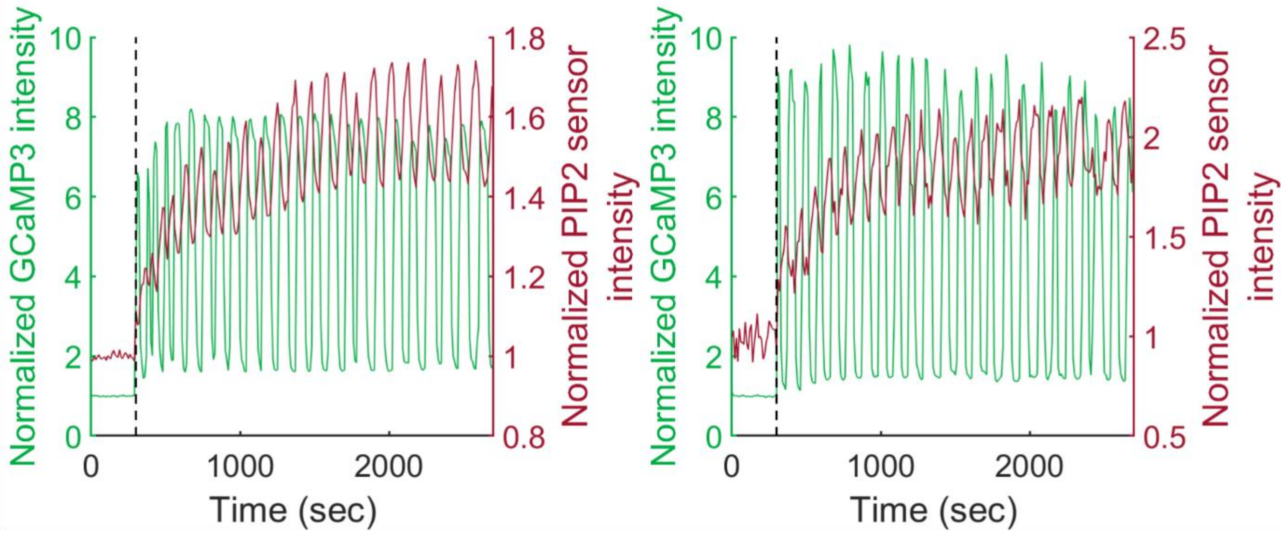


Figure S1. Exemplary cells with constant oscillation from co-imaging of Ca2+ and PIP2.

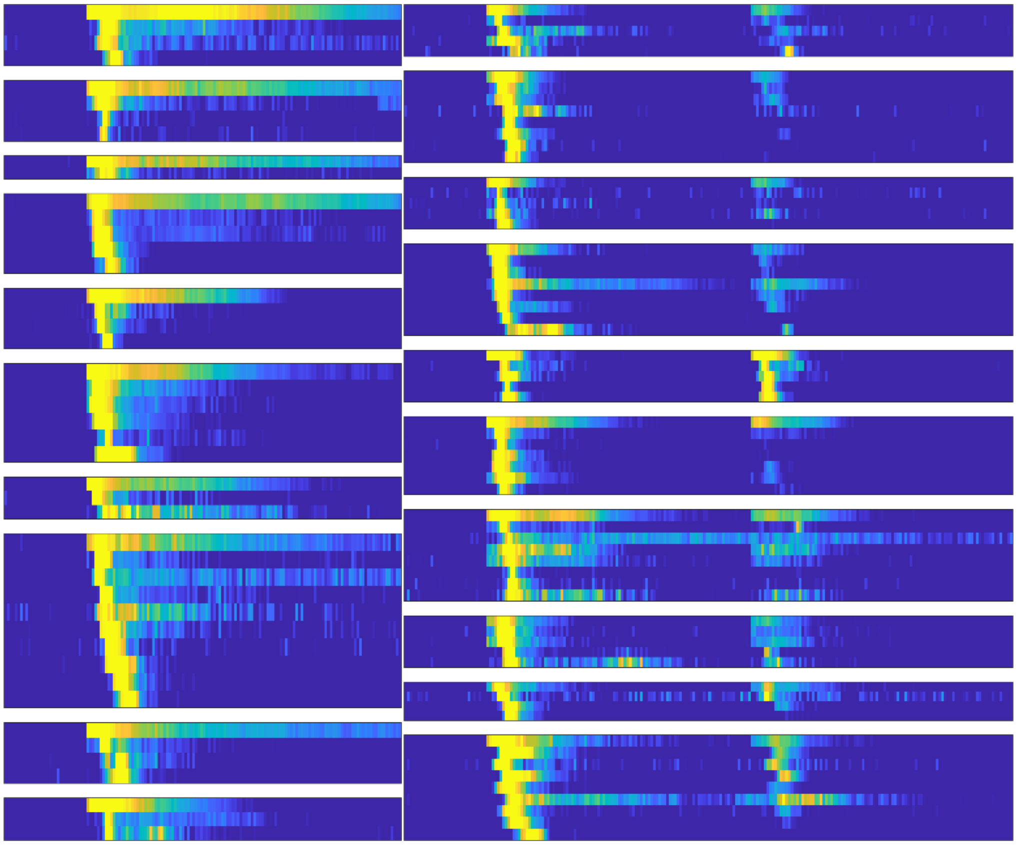

Figure S2. Representative $Ca^{2+}$ traces from hub and follower cells from ‘pattern 1’ (left) or ‘pattern 2’ (right) stimulation. Each individual subgraph shows the $Ca^{2+}$ time series from one hub cell and its follower cells. The $Ca^{2+}$ time series of hub cell is shown at the top of each subgraph and that of follower cells are sorted based on the time point of reaching peak.

| Abbreviation | Meaning | Default Value |
|---|---|---|
| $v_1$ | Voltage of half opening for VDCC | -20 mV |
| $v_2$ | Voltage sensitivity of VDCC | 24 mV |
| $v_3$ | Voltage of half opening for Kv | -40 mV |
| $v_4$ | Voltage sensitivity of Kv | 18.4 mV |
| $g_{Kv}$ | Conductance of Kv channel | 180 pS |
| $g_{Ca}$ | Conductance of VDCC | 1000 pS |
| $g_{KATP}$ | Conductance of KATP | 4000 pS |
| $E_K$ | Nernst potential of $K^+$ | -75 mV |
| $E_{Ca}$ | Nernst potential of $Ca^{2+}$ | 25 mV |
| $C_m$ | Membrane capacity | 5300 pf |
| $K_{ATP}$ | Dissociation constant of ATP-KATP complex | 0.012 mM |
| $K_{ADP}$ | Dissociation constant of ADP-KATP complex | 0.45 mM |
| R | Glucose-sensitive mitochondrial rate parameter | 0.9 |
| $R_1$ | $Ca^{2+}$-sensitivity of mitochondrial proton-motive force | 0.35 mM |
| $k_{mito}$ | Rate constant for ATP hydrolysis | $5*10^{-5}$ $ms^{-1}$ |
| $A_{total}$ | Total concentration of ATP and ADP | 1 mM |
| fi | Fraction of free $Ca^{2+}$ | 0.01 |
| α | Factor for converting $Ca^{2+}$ current to concentration change | $4.5*10^{-6}$ uM*(fA*ms)$^{-1}$ |
| $k_{CaL}$ | Rate of $Ca^{2+}$ removal from cytosol | 0.15 $ms^{-1}$ |

Table S1: Parameters used in mathematical modeling.
